# Subjective, not objective, socialness drives activity in the brain’s third visual pathway

**DOI:** 10.64898/2026.05.28.728411

**Authors:** Rekha S. Varrier, Qi Liang, Tory G. Benson, Peter J. Molfese, Emily S. Finn

## Abstract

Recent work, including the third visual stream hypothesis, frames social perception as a largely automatic visual process. Simple animations of shapes engaged in “social” behaviors such as chasing have long been used to study social perception; however, both behavioral and neural responses are typically modeled against discrete, experimenter-assigned stimulus labels. This approach is insufficient, given that perceptions of such stimuli vary not only between people but also across trials. Here, we directly compared the neural responses to objective (stimulus-based) and subjective (response-based) socialness in an fMRI study in which the degree of chase between two agents was varied algorithmically. Participants (n=24) rated their perception of a chase on a continuous scale. Subjective ratings better predicted activity in visuo-social brain regions, predominantly in area MT, even after controlling for optic flow. Our findings underscore the visual nature of social cognition and highlight the importance of analyses sensitive to idiosyncrasies in perception.

## Introduction

Humans are an obligate social species; our survival depends on accurately recognizing and categorizing social information. We might therefore expect these evolutionary pressures to have given rise to specialized cognitive mechanisms for processing social interactions, and neural substrates to support those mechanisms ^1–4^.

Fully processing and understanding social interactions involves several steps, including identifying animate agents, deciding whether they are engaged in goal-directed behavior and whether that goal-directed behavior is *social* in nature, and attributing beliefs or motivations to explain that behavior; these steps likely rely on partially distinct but interacting brain networks ^5,6^. Based on classic work, the earliest of these steps – the identification of animate agents and social interactions between them—was thought to activate both the lateral occipital cortex ^7^ and regions in the action observation network like the posterior superior temporal sulcus (pSTS; Castelli et al., 2000; Gao et al., 2012; Schultz et al., 2005) and the parietal lobe ^11,12^. While high-level mentalizing is thought to recruit canonical “social brain” regions such as the medial prefrontal cortex and temporo-parietal cortex ^13^, recent work has suggests that the earliest steps in detecting social interactions are largely automatic and visual in nature ^14,15^, and that there likely exists a third visual stream beginning in primary visual cortex and proceeding laterally along the occipitotemporal cortex to facilitate rapid, perception-based recognition of social information ^16^.

Even when social interaction recognition feels automatic, almost effortless, there is some variability both across and within individuals in if and how information is classified as social, as noted by even the earliest studies ^17,18^. It is important to respect this variability for two reasons. First, we lack an objective, “ground-truth” definition for what makes information social: real-life interactions are highly culture and context-dependent, so to some extent, “socialness” only exists in the subjective sense ^19^. While certain stimuli—such as a tennis match or a dyadic conversation—very obviously involve a social interaction, other types of stimuli can evoke strong percepts of “socialness” in some individuals or contexts but not others. Consider determining if someone is trying to catch your gaze from across a room, or phenomena such as pareidolia (in which we detect illusory faces in inanimate objects ^20^) or the Heider-Simmel effect, in which animated videos of simple geometric shapes can evoke rich social meaning from otherwise unlifelike scenarios ^17^ . Rather than dismissing these as edge cases, we can harness them to more fully characterize the properties and limits of the human cognitive mechanism for social signal detection ^21^. Second, deviations from the “typical” or average percept can be related to mental illness and other conditions: consider hypo-mentalizing in autism or hyper-mentalizing in some psychotic disorders ^22–26^.

Here, we capitalize on a class of social stimuli that are inherently ambiguous—namely, Heider-Simmel-style shape animations—to rigorously test the hypothesis that “socialness is in the eye of the beholder”. Some previous work, including our own, has found stable inter-individual differences in socio-perceptual thresholds ^27^ and that subjective percepts are superior to objective (experimenter-defined) labels when explaining brain activity ^12,21,28^. Here, we extend this work using carefully controlled stimuli and a robust within-subject approach to show that regions of the third visual stream more closely track levels of perceived socialness than objective evidence for socialness. We used an established approach ^10,29^ to parametrically vary the degree of movement contingency between two agents (objective social evidence) and elicit a continuous rating of ‘socialness’ from each participant on each trial (subjective social reports). In both whole-brain (data-driven) and region-of-interest (hypothesis-driven) analyses, we show that univariate activation magnitude in regions of the third visual stream, including middle temporal area (MT), is sensitive to subjective reports over objective evidence, suggesting that the presence of information ultimately classified as social modulates activity early in the perceptual-cognitive hierarchy.

## Results

### Parametric animations evoke variation in socialness judgments

In this study, we parametrically varied a mid-level motion attribute known to modulate socialness judgments in 6-second animations involving two agents represented as dots, one black and one gray. In these animations, one agent (“predator”) is programmed to follow the other (“prey”; agents’ colors randomized across trials) at varying levels of *chase directness*: 0 (no following), 0.167, 0.33, 0.5, 0.66, 0.833 and 1 (very clear following). Further, we included a control condition in which the predator followed an invisible prey at an arbitrary location, whose motion contingencies were mimicked by the second agent (or fake prey) on screen (for details, see Methods). Social perception (or “socialness”) was measured in this study as the rating on a continuous scale ranging from “Random” to “Follow”.

In line with past work ^10,27,29^, we observed that socialness scales with *chase directness* in the true chase condition, but not the invisible-chase control condition (Fig. 1). Further, to verify that the detected pursuit was indeed the intended one, we had a secondary attention-check task, namely, to identify the predator agent (by naming its color). As with the subjective perceived socialness, objective predator detection accuracy also increases with *chase directness* (Fig. S1). While the overall trends are largely preserved in each individual participant (Fig. S2), there was also considerable variability in socialness judgments both within and across participants, which can be observed by the spread of responses at each *chase directness* level (individual data points) and overall shape of the curve, respectively (Fig. 1b, S2).

**Fig. 1:**
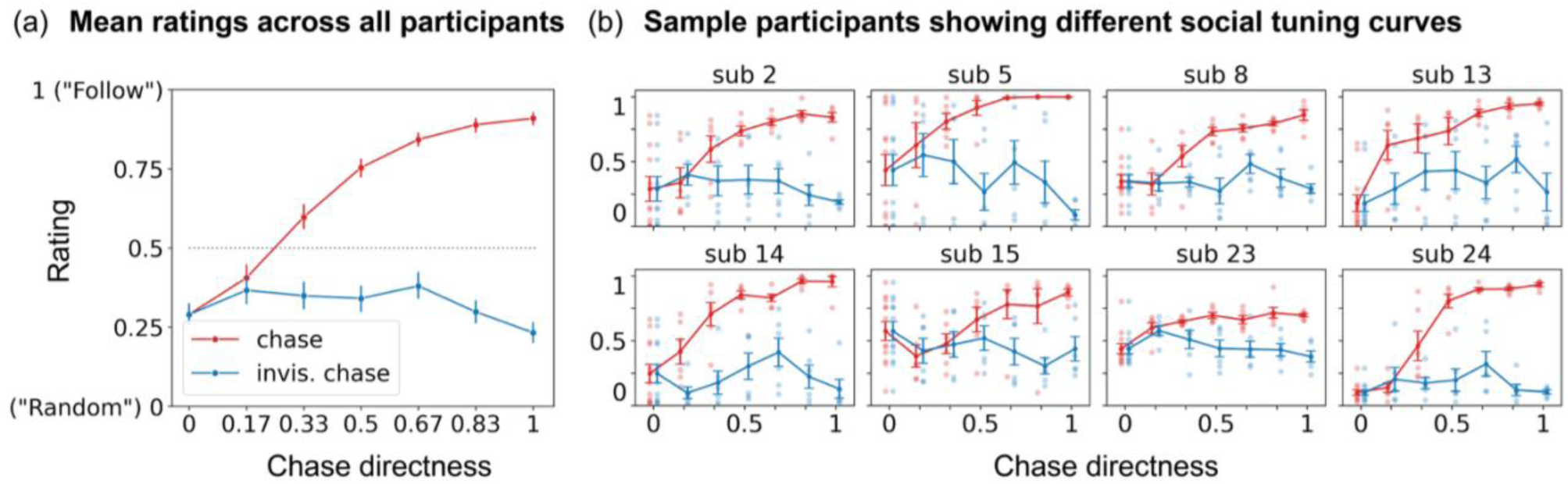
Behavioral data. (a) Mean data across all participants, and (b) from 8 sample participants (out of 24 total). (a) As chase directness increases (X-axis), “follow” perception (Y-axis) increases in the true chase animations (red), but not the control animations (blue) in all participants (full set of participants in Supplementary S2a). Error bars show 95% confidence intervals. (b) The difference in shape and position of each curve (i.e., line connecting mean ratings at each chase directness level) across participants suggests the presence of individual differences in socialness judgments that exist atop group-level similarities. The dots within each panel (representing trial-level ratings) and corresponding error bars (representing standard errors of the mean) show that within individuals, socialness ratings can vary across trials at the same chase directness level.

### Both visual and association areas respond to perceived levels of socialness

Having verified that our animation stimuli evoked variable amounts of perceived socialness, we next sought to identify brain regions where activity levels tracked subjective ratings of socialness.

To this end, using data from all trials, we modeled trial-wise activation magnitude as a function of the participants’ own socialness rating given at the end of each trial. A whole-brain analysis revealed several regions where activity increased as perceived socialness increased (Fig. 2). These included primary visual and visual association regions like the left primary visual cortex, the bilateral MT and the fusiform face area (FFA); parietal regions like the right supramarginal gyrus, superior parietal lobule (SPL), and intraparietal sulcus (IPS); parts of the postcentral gyrus and left medial parietal lobe; frontal regions like the precentral gyrus, supplementary motor area (SMA) and inferior frontal gyrus (IFG); and medial cerebellum. This analysis also revealed some areas where activity *decreased* as perceived socialness increased, including the left superior temporal gyrus (STG), early visual cortex, ventral visual cortex and parts of the supramarginal gyrus and anterior cingulate cortex (ACC).

**Fig. 2:**
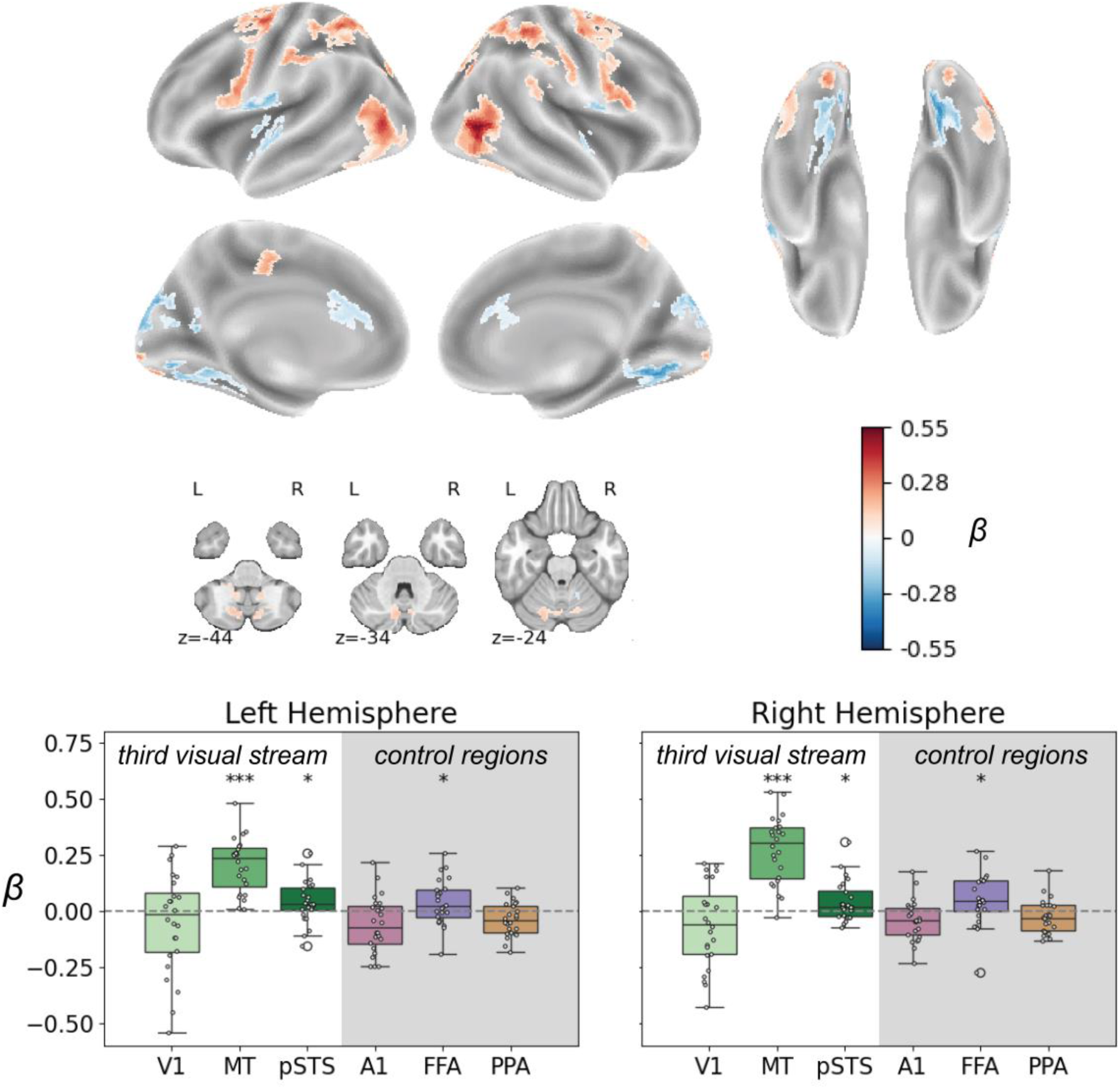
Brain regions that respond to subjective ratings of socialness. Brain surface and cerebellar views show results from a whole-brain voxel-wise analysis (map thresholded at q < .05, corrected for multiple comparisons using the false discovery rate, plus a nominal cluster size threshold ≥ 30 voxels). Boxplots in the lower row show results from a region-of-interest (ROI)-based analysis (one-sampled t-tests against 0; one-tailed) focusing on regions of the third visual stream as well as control regions, separated by hemisphere. ROIs definitions are shown in Supplementary Fig. S3. *** = p < .001, * = p < .05.

In a region-of-interest (ROI) analysis in which we focused on ROIs from the third visual stream, we find that two of them—the bilateral MT and posterior superior temporal sulcus (pSTS)—as well as the ventral visual control region FFA show higher activity in response to higher levels of socialness, whereas the primary visual cortex (V1) and the other two control regions—the primary auditory cortex (A1) and parahippocampal place area (PPA)—do not (Fig. 2b).

Importantly, we find that the response of MT, a region typically known as a low-level motion perception region, to the degree of perceived socialness holds true even after regressing out the effect of the total optic flow, a measure of the total amount of motion within each animation (see supplementary Fig. S4 and **Methods** sub-section **Optic flow**).

### Subjective socialness ratings explain neural activity better than objective motion attributes

The previous analysis suggests that several brain regions track levels of perceived socialness. However, subjective socialness ratings covaried substantially with objective evidence for socialness (i.e., *chase directness*). Which predictor—subjective rating or objective evidence—better explains trial-to-trial variance in activation magnitude?

To answer this, we directly contrasted outputs from models using subjective (Fig. 3a) versus objective (Fig. 3b) ratings of socialness. (This analysis was limited to chase trials, since the objective measure of *chase directness* was not meaningful in invisible-chase [control] trials.) In a whole-brain analysis, subjective ratings explained more activity especially in the MT at an uncorrected threshold (*p* < .001 uncorrected, Fig. 3c). Overall, subjective ratings were a better predictor of activity in more brain regions than objective *chase directness* (Fig. 3 a-c, supplementary Fig. S6 and S7). Further, ROI analyses confirmed that both left and right MT were more responsive to subjective than objective ratings (Fig. 3d).

**Fig. 3:**
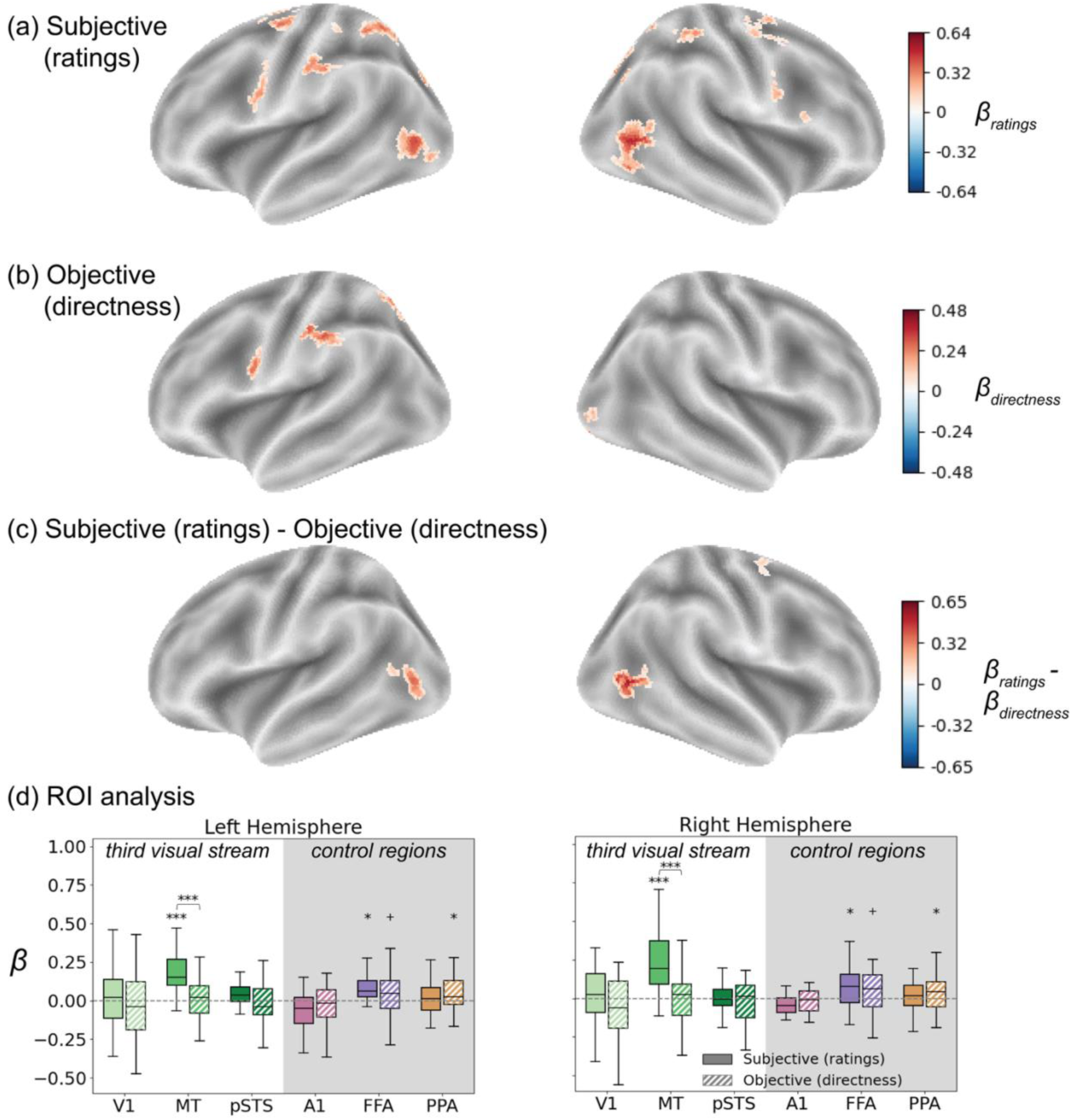
Brain activity in response to (a) subjective ratings, (b) objective directness, and (c) ratings – directness (whole-brain voxel-wise analysis thresholded at p < .001, uncorrected for multiple comparisons, with a nominal cluster size ≥ 30 voxels). (d) Region-of-interest (ROI) analyses showing one-sample t-tests against zero for both subjective and objective predictors, as well as paired t-tests directly comparing subjective versus objective predictors. *** = p < .001, * = p < .05, + = p < .1.

## Discussion

Using simple animations with varying evidence for a social interaction, we found that activity in MT, a key region in the third visual stream, tracks the observer’s perceived levels of socialness, even when controlling for optic flow and other objective stimulus attributes. This result adds to a growing body of work suggesting that key aspects of conscious social perception rely automatic, bottom-up visual processes that take place in brain regions early in the perceptual-cognitive hierarchy.

While visual region MT has long been known to be sensitive to motion in general, its preferential role in processing social motion has only begun to be appreciated recently. Even though some past work had alluded to a role of area MT in biological motion processing—for example, people on the autism spectrum showed less activity in MT when identifying non-social biological motion even when behavioral accuracy did not decrease ^30^, and social video clips elicited higher activation in the right hemisphere than non-social clips ^31^—traditionally, it has not been widely considered a relevant area for social perception more broadly. Moreover, studies have largely not been able to tease apart sensitivity to objective (i.e., stimulus-computable) motion attributes from sensitivity to subjective (i.e., observer-reported) percepts of “socialness”, since most past work has used handcrafted animations intended to appear either “obviously social” or “obviously non-social”, then modeled brain activity with respect to these labels without accounting for participants’ actual percepts. In a previous study using geometric shape animations ^21^, we found that activity in many brain areas, including MT, is better explained by observer reports than the intended experimenter labels, previewing the results reported here. Still, our earlier study, like many of its predecessors, relied on a small number of experimenter-crafted animations that also varied along uncontrolled dimensions such as number of agents, shape, color, speed, and the presence of inanimate objects. Here, we more rigorously test the hypothesis that activity in visual regions tracks perceived socialness by using parametrically generated stimuli that vary along a single dimension of interest, namely, the degree of pursuit-like motion contingency between two agents (operationalized as *chase directness*), while keeping all other visual attributes as constant as possible. (While some variance in optic flow across levels of *chase directness* was unavoidable in our stimulus-generation algorithm, results persisted when controlling for this additional variable.) Leveraging deliberately ambiguous stimuli—i.e., those that yield high variability in reported percepts—we show that MT tracks *perceived* socialness above and beyond *objective evidence* for socialness, positioning this region as a key early node in consciously processing and evaluating potential social information. We also note the possible overlap between MT and the extra-striate body area, an area that encodes relative dyadic body configurations ^32^ and may also be part of the third visual stream ^16^.

Past work using social animations reported pSTS as central to animacy and social perception ^9,10,33,34^. In our study, while we saw some socialness-related activity in pSTS (Fig. 2), it was overall weaker than activity in the more upstream region, MT. This difference could be due to the simplicity of our stimuli, which were overall quite short in duration and simple in the interaction they depicted – there were no physical barriers for agents to navigate, ^34^ nor interactions evoking clear valence like helping vs. hindering ^33^, nor a predator with changing intentions ^9^. This likely means that making judgments about our stimuli recruited less mentalizing and more automatic, bottom-up perception. Hence, unlike previous Heider-Simmel-style animations, our animations likely did not spontaneously prompt participants to mentalize about the *causes* behind any possible social interaction (i.e., the dots’ motivations or beliefs), and percepts were likely driven by the motion patterns themselves. Recent work suggests that the pSTS, along with other canonically social regions like the temporo-parietal junction (TPJ), may perform both bottom-up and higher-level inferential computations at different timescales ^35^; it is likely that longer and more complex animations would have elicited more activity in pSTS and later regions.

In addition to areas of the third visual stream (MT and pSTS), many other brain regions tracked subjective socialness (Fig. 2), many of which have been associated with social scene navigation and/or action perception. For instance, the superior parietal lobule is both part of the action observation network ^36^ and has been reported to be involved in navigating visual scenes ^37^, whereas the inferior parietal sulcus has been reported to encode levels of animacy ^12^. Fusiform face area is sensitive to biological motion ^38^, animate and rational agents (compared to inanimate controls ^39^) and social cognition ^40^. The frontal regions postcentral gyrus, supplementary motor area and inferior frontal gyrus have been linked to using social cues to make decisions ^41^. Some regions also showed lower activity with greater socialness, such as the right medial occipital and left superior temporal areas; these areas could be reflecting the higher attentional demands on the visual system due to noisier motion and the need to focus on two agents instead of one social dyad ^42^ as well as longer decision-making times when watching the less social stimuli. However, in the direct contrast between subjective socialness ratings and the objective stimulus attribute, we found subjective ratings to show relatively stronger responses in area MT alone.

Our strong neural evidence for the idea that “socialness is in the eye of the beholder” has important implications for computational modeling and artificial intelligence. Existing work has sought to describe in formal terms the spatial and temporal motion contingencies between agents that give rise to social percepts *on average* across people ^29,43–46^, and to model the mechanisms by which people might arrive at these judgments ^47–52^. This work has made exciting strides in our algorithmic understanding of *if* and *how* social interactions are recognized. However, analogous to the experimenter-derived labels used in most past behavioral and neuroimaging work described above, most of these modeling efforts assign training labels by taking the consensus from a set of human judgments, ignoring the often considerable variability in those judgments. Ultimately, because these models are only as good as the ground truth used to train them, understanding heterogeneity in whether and how social interactions are recognized will be essential to a complete explanation of social perception.

We acknowledge some limitations to the work presented here. First, for the region-of-interest analyses, we did not have functional localizers in individual participants; instead, we relied on standard anatomical maps and meta-analytical functional maps. This could have led to some false negatives (missing relevant participant-specific voxels) and false positives (more noisy voxels). Second, there could have been possible confounding effects of animacy percepts on socialness ratings, since both animacy and socialness could have decreased as *chase directness* increased. This can potentially be addressed by using a cover story to set a context so that the agents “mean” something, for example, label agents as wolf and sheep as in Gao et al. ^29^ or as children in a park like in our previous study ^27^. Lastly, self-report itself is prone to reporting biases, which can be addressed in future work using eye-tracking and physiological data to complement subjective reporting.

In conclusion, social perception is at the same time a high-level and low-level process – high-level because it results in complex percepts even with such simple stimuli, and low-level because as we have shown here, for simple visual contingencies, visual areas play a critical role in inducing social percepts.

## Star Methods

### EXPERIMENTAL MODEL AND STUDY PARTICIPANT DETAILS

#### Participants

Thirty-five individuals took part in the study, out of which twenty-four met criteria for our final analyses (age: mean ± SD = 27.42 ± 5.32 years old; gender: 11 female, 10 male, 2 non-binary, 1 unreported). Details of data exclusion are described in the *Data pre-processing* section. Participants were recruited by advertisements within Dartmouth College and the surrounding community. Prior to in-person testing, potential participants underwent eligibility screening via an online survey. Eligibility criteria were as follows: 18-45 years old, fluent in English, no claustrophobia, no unremovable metal in body, normal or corrected-to-normal vision (with contact lenses), no serious psychiatric illnesses, no history of serious head injuries/concussions, weigh under 300lb, are right-handed or ambidextrous, are willing to take part in two sessions spaced 3-8 weeks apart, and are willing to continue even after being given information in writing about how MRI studies are conducted and what will be expected of them (to lie still for up to 1.5 hours), along with a link to a short clip with scanner noises (https://www.youtube.com/watch?v=hvXoHU9Cexk). All procedures were approved by the Committee for the Protection of Human Subjects at Dartmouth College.

### METHOD DETAILS

#### Stimuli

##### Motion attribute: Chase directness

We generated animation stimuli using the first iteration of the custom Javascript-based software called *psyanim* developed in our lab (https://github.com/thefinnlab/psyanim). Each animation lasted 6 seconds and consisted of two agents, represented by dots (one gray and one black), moving around the screen. Stimulus generation – particularly the motion attribute used to parametrically generate stimuli in our study – was identical to that described for the social detection study in our recent behavioral paper ^27^. Both of our studies were inspired by past work by Gao et al. ^29^, who showed that the perception of a chase decreased with the parameter called “chase subtlety”, which controls how directly (or subtly) a predator agent moved towards a prey agent.

For the current study, we created 168 animation stimuli in total. In 72 of the animations (“chase” condition), one agent (chosen randomly) assumes the role of predator and the other assumes the role of the prey. The predator was programmed to follow the prey at a given level of *chase directness* (see below), and the prey was programmed to evade the predator. In 72 of the remaining animations (the “invisible-chase” condition), the predator was programmed to chase a prey, also at a given level of *chase directness*, except that the true prey was invisible and at a random location on the screen. Similar to Gao et al (2009) and our recent paper, the second visible agent (“fake prey”) mimicked the invisible prey. By “mimic”, we mean that the fake prey copied the true prey’s trajectory, but with a 180° rotation (i.e., if the invisible prey moves down and to the right, the mimicking agent moves up and to the left). This condition was to meant serve as a control where the visible agents preserved temporal relationship of the chase condition, but their spatial relationship was incongruent (in the true chase, the spatial and temporal relationships were congruent in the more direct chases), and hence the percept that one agent was following the other agent would be destroyed.

To vary the degree of perceived social interaction (i.e., the chase), we parametrically varied the motion attribute *chase directness*. *Chase directness* was defined as 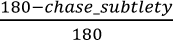, where *chase_subtlety*, first defined by Gao et al. (2009), is the angle by which a predator could deviate from a perfect heat-seeking path between itself and the prey at each time step. Lower (higher) the *chase directness (chase_subtlety)*, less directly (more subtly) the predator follows the prey. Thus, at *chase_subtlety* = 0° (*chase directness* = 1), the predator would directly pursue the prey at each timestep, so that it would be clear to observers that the predator was following the prey, whereas at *chase_subtlety* = 90° (*chase directness* = 0.5), the predator goes sometimes in the direction of the prey, and sometimes in a random direction between the direct path towards the prey and 90°clock-wise or counter-clockwise, resulting in a noisy chase. Up to *chase_subtlety* = 90° (*chase directness* = 0.5), the predator would be only moving in the general direction of the prey, but at *chase_subtlety* > 90° (*chase directnesss* < 0.5), the predator would sometimes move towards the prey and sometimes away from it. In the current study, we generated stimuli at 6 levels engaging in the predator-prey interaction: *chase_directness* 0.167, 0.33, 0.5, 0.667, 0.833, 1 (*chase*_*subtlety*: 0°, 30°, 60°, 90°, 120°, 150°). Stimuli for *chase_subtlety* = 180° (*chase directness* = 0) was operationalized as both the predator and the prey “wandering” around the screen.

##### Stimulus generation

Each animation was 6 seconds long (frame rate: 30 frames/s; 180 frames in total) and consisted of two circular agents (one black, one grey, 40px diameter). The initial positions of the agents in pixels were [350, 450] and [850, 450] for the agent in the left and right, respectively. The predator and prey agents’ colors and initial positions were varied randomly across conditions. For the first second, the two agents remained static in their starting position on the left and right side of the center of a 1200px x 900px screen. After the 1 second static period, both agents moved around the screen as programmed until the end of the animation. The animations can be downloaded here: https://github.com/thefinnlab/psyanim1_fmri.

As with our recent work ^27^, stimuli were generated programmatically; all motion attributes except the *chase_subtlety* were kept constant across stimuli. Based on the *chase_subtlety*, on each animation frame, the predator’s motion direction was selected from the range [-*chase_subtlety*, *chase_subtlety*]. Selection of the direction was from an almost-uniform distribution, which was accomplished by choosing a random value between 0 and 1 which then mapped on to *chase_subtlety* via a 4-parameter (cubic) Bezier ease-out function (from the npm-dependency *bezier-easing* version 2.1.0) with the parameters x1: 0.25, y1: 0.5, x2: 0.5, y2: 0.75. Since these parameters generate a curve that starts steeper and ends flatter compared to a linear function, in effect, this way of choosing motion direction on each frame reduce the odds of sudden fluctuations or jitteriness in the motion direction of the predator which can occur by chance with the selection of extreme values compared to intermediate values (e.g., this function reduces the probability of the predator moving upwards in one frame and downwards in the frame immediately after that). There were 12 animations for each of the 6 levels of *chase directness* both in the “chase” and in the “invisible-chase” conditions. At each level of *chase directness*, predator identity (gray or black) and starting location (left or right) were fully counterbalanced.

All other attributes were kept constant across all animations (these values were determined after extensive piloting): (i) The maximum speed of the predator (50px/s) was set to a value slightly smaller than that of the prey speed (60px/s) – the difference was to ensure that the predator never actually caught up to the prey in the animation; (ii) The predator would almost always chase the prey (if it is within 1000px, parameter *boredom distance*), but the prey would flee only if the predator was within a 300px radius (*safety distance*); (iii) the prey moved with a *flee_subtlety* of 60° (defined by how much the prey could deviate from a perfect fleeing path – i.e., instead of always moving in the exact opposite direction of the predator, the movement pattern was made slightly noisy). This last parameter (*flee_subtlety)* was implemented to prevent the fleeing motion of the prey, as opposed to the overall interaction between the agents, driving percepts. Both agents were also programmed to avoid edges and corners of the screen.

For animations with *chase_directness* = 0 (*chase_subtlety* = 180°), both agents “wandered” (i.e., moved without any specific pre-assigned goal) on the screen, with maximum speeds matching those of the true predator and prey, respectively, as described above.

The final stimuli were generated programmatically using *psyanim* and quality-controlled manually with the help of research assistants. All animations were pre-generated and displayed as video clips (*webm* format) on the screen at experiment run time.

#### Task design

Each participant completed 84 trials (6 trials each of each of the 6 *chase_directness* levels (0.167 to 1) of the chase and invisible-chase stimuli, and 12 wander trials in lieu of *chase_directness* = 0) – the exact trajectories never repeated, each trial presented a unique stimulus. Stimuli were pseudo-randomized such that all levels of *chase_directness* and condition (chase, invisible-chase and wander) were equally present within each quarter, i.e., each set of 21 trials (of this, 9 each were chase and invisible chase (6 unique *chase_directness* levels and 3 randomly selected; the randomly selected *chase_directness* levels were counter-balanced between quarters 1 and 2, and 3 and 4) and 3 represented chase_directness 0). Participants watched each animation and (i) rated the socialness of their percept on a continuous scale ranging from “Random” to “Follow” (henceforth often referred to as “socialness ratings”) and (ii) identified the predator agent by color (two-alternative forced choice). They were told to guess when unsure. To make responses, participants used a mouse with a track ball that they could rotate to move around the screen as needed and click to finalize decisions. One participant (sub 1) had pressed the wrong button for the second question (they clicked on the agent that was *not* the predator), and to correct for this, we reverse-coded their responses.

#### Summary of pilot behavioral experiments

Prior to the fMRI study, we first tested the efficacy of these stimuli in eliciting variable social percepts in a series of behavioral experiments conducted online. Data from 3 batches of 200 participants each were collected online using jsPsych ^53^ on Prolific (www.prolific.com). In the earliest pilots, we collected two batches of data where in one group, participants rated stimuli on a continuous scale ranging from Random ↔ Follow and in the other group, they rated it from Random ↔ Social. In the latter group, we later also collected free-text responses for a subset of trials in which participants were additionally asked to describe the scene using the prompt “*Write down, in a sentence or two, what happened between the shapes (if anything)*” to assess whether social interactions indeed referred to chase-like interactions as intended. All pilot data confirmed that as *chase_directness* increased, so did the perception of socialness, whether measured using the more specific “Follow” label or more general “Social” label. Both versions also showed notable across-trial variability in ratings even at the same *chase_directness* value (a desired feature here) and across-participant differences in terms of the shapes of the psychophysical curve. The text analyses also showed an increase in the reporting of chase-like behaviors as the *chase directness* attribute increased, confirming that “Follow” was a reasonable choice. We chose the more specific label for our main experiment since it was overall more reliable.

#### fMRI experiment

Participants were briefed in detail about the experiment outside the scanner and reminded of the instructions once they were in the scanner. The experiment was programmed using PsychoPy3’s Builder interface ^54^.

##### Data acquisition

Data was collected on a Siemens Magnetom 3T Prisma scanner. Each session started with a quick scan or an anatomical scout (14s). Next, there was a T1 scan (sequence: MPRAGE) where participants were told to either look at a fixation cross or keep their eyes closed. Voxel size: 0.9mm x 0.9mm x 0.9mm, TR = 2.3s, TE = 2.32ms, flip angle = 8°. The T1 scan lasted 5 minutes 21 seconds. Following this, there was a field map (20s) before the main functional phase. For functional imaging, we used EPI (echo-planar imaging) sequences with the following parameters: TR=1.057s, 52 slices, voxel dims: 2.7mm x 2.7 mm x 2.7mm, multi-echo sequence (TE1 to 3: 1.42ms, 34.42ms, 54.64ms, respectively), flip angle 59°, phase encoding direction A – P. Each run started after receiving the fMRI trigger. The beginning and end of each run were padded with a 10 sec fixation block. Each participant performed 84 trials split into in 7 runs (12 trials/run).

Each trial started with a pre-stimulus fixation window of 5 seconds (± 2 seconds jitter) followed by the animation stimulus (6 seconds). After this, there was a post-stimulus fixation period of 2 seconds (± 1 second jitter), followed by a response screen (described in detail in **Task design**). Each run varied in length, since participants could complete individual trials at their own pace (with an upper limit of 10 seconds on the response screen). To accommodate the slowest possible participant, we set the EPI sequences in each run to 365 scan volumes (maximum possible duration including the pre-scan delay: 6 minutes 46 seconds). Each run was stopped manually when the task was completed.

Additional measures were collected. Participants came back for a second scan session that was identical to the first session described above, except that at the end of session 2, we additionally acquired an 8 min resting state scan from 22 of the 24 participants. Participants were asked to fixate on a cross at the center of the screen during this run (same EPI sequence as above). At the end of a scanning session, participants completed a series of trait questionnaires before being debriefed. In addition to fMRI, eye-tracking data were collected in the scanner using an MR-compatible EyeLink 3 system from SR Research. Data from session 2, eye-tracking and the traits are not analyzed as part of the current paper; here, we only report our findings from fMRI data collected during session 1.

##### Data pre-processing

The raw DICOM images were first converted to the BIDS format using the HeuDiConv (a heuristic-centric DICOM converter; https://github.com/nipy/heudiconv). Skull-stripping of the T1w image was performed using Freesurfer (https://surfer.nmr.mgh.harvard.edu/). Pre-processing was done using AFNI (https://afni.nimh.nih.gov/) using the following steps by customizing the afni_proc.py file: (i) aligning the 2^nd^ echo (from the multi-echo scans) to the skull-stripped T1w image, (ii) aligning to the MNI template image MNI152_T1_2009c+tlrc, (iii) motion-correction and alignment to the EPI volume with the fewest outlier voxels in each run, (iv) distortion-correcting the field using the field maps collected prior to the functional scans, (v) masking the voxels using the EPI scans, (vi) combining the multi-echo images using *tedana* (https://tedana.readthedocs.io/en/stable/), (vii) applying a spatial smoothing of 4mm, and (viii) rescaling timeseries to a mean of 100. Additionally, for subsequent quality-checks, (i) a basic GLM (stimulus > rest) was run and the results from the t- and F-statistics plotted, and (ii) censoring thresholds were set – to remove scans (TRs) where the estimated head motion > 0.2mm and scans where more than 5% of voxels are statistical outliers. We used AFNI’s own recommendations to manually quality-check (QC) and exclude participants ^55^. We excluded 8 participants who showed excessive motion – specifically, we excluded (i) participants who showed a head motion either of more than 0.1mm on average during stimulus periods in the task (QC param *ave mot per sresp*) or across all TRs in the study (QC param *ave censored motion*) and (ii) participants with a lot of “censored” data, i.e., more than 10% TRs flagged as bad based on the exclusion criteria described in point (i) or missing voxels due to motion. Two more participants had already been excluded prior to pre-processing – one participant’s brain showed up tissue damage due to a head injury from early childhood during the scan (unknown to us at the time of recruitment), and a second participant performed poorly in the behavioral tasks (both rating the socialness and identifying the predator; confirmed as fatigue during the post-scan debriefing). The quality-check led us to exclude 10 participants in total, leaving us with 24 participants with good-quality MRI data.

### QUANTIFICATION AND STATISTICAL ANALYSIS

First (subject)-level and second (group)-level statistical analyses were also performed using AFNI.

#### General linear model (GLM)

We performed separate analyses to answer two questions: first, how the brain responds to parametric levels of subjective socialness, and second, how the model that fits brain activity to subjective ratings compares to the model that fits objective ratings.

##### Analysis 1: Neural response to subjective socialness

First, we aimed to identify voxels that respond to levels of perceived (subjective) socialness. For each subject, using data from all trials (both chase and invisible-chase), we modeled the BOLD timecourse as a function of socialness ratings using amplitude-modulated block regressors (where the modulation came the participant’s own rating collected at the end of each trial) of duration 6 seconds (matching the onset time and duration of each animation), using the 3dDeconvolve function in AFNI. We used an additional amplitude-modulated regressor for optic flow (defined as the overall motion contained in the 6 second animation, see details later) and an unmodulated regressor of magnitude 1 for the response window whose onset is the beginning of the response window and whose duration matches the response time of each trial). Both socialness ratings and optic flow were normalized (rescaled to 0-1) before fitting. The 6 motion regressors (3 translational and 3 rotational dimensions) estimated in the motion correction step described above were entered as additional covariates, as well as a second-order polynomial baseline model for baseline and temporal effects – this adds a constant term (mean activity per voxel per run), a linear drift term and a quadratic drift term. After defining the model, voxel-wise estimates were derived using the 3dREMLfit which fits the model after also factoring in temporal auto-correlation. We performed second-level (group) analysis as a one-sample t-test using the 3dttest++ function in AFNI, giving us group-level z-scores for each voxel.

##### Analysis 2: Comparison between neural responses to subjective and objective socialness measures

For this analysis, we excluded the invisible-chase animations, since the term *chase directness* was a meaningful contrast to subjective ratings only for the true chases. Because subjective (ratings) and objective (*chase directness*) socialness were substantially collinear, rather than include them both as predictor terms in the same model, we performed two single-subject GLM models: one where the trial-by-trial amplitude modulation came from the subjective ratings (similar to analysis 1, but excluding the invisible-chase trials), and a second where the trial-by-trial amplitude modulation came from the objective motion attribute *chase directness*. Besides these, each model additionally contained optic flow and the response window similar to model 1. Since invisible-chase trials were excluded from this analysis, our model definition replaced these with 0s in the block regressors. We then compared the resulting maps at the second- (group) level using a paired t-test using the 3dttest++ function in AFNI, giving us group-level z-scores for each voxel.

#### Whole-brain and ROI-based analyses

With the voxel-wise z-score maps, we performed our analyses at two-levels. First, in a whole-brain analysis, we estimated the statistical significance per voxel with multiple-comparison correction (false discovery rate) across the grey matter voxels in the brain, and created whole-brain activation maps using *nilearn.* Second, we defined regions of interest (ROIs) along the main regions that form the third visual stream comprising the primary visual cortex (V1), MT and the posterior part of the superior temporal sulcus or pSTS ^16^. We additionally defined three control regions where we expect little to no activity for social motion: the primary auditory cortex (A1), fusiform face area (FFA) and parahippocampal place area (PPA). Because we did not collect individual functional localizers from our participants, we used a combination of independent functional and anatomical maps to define our ROIs. Functional masks from meta-analyses were selected using the z-scores (thresholded at *q* < .01 and FDR-corrected for multiple comparisons) from the association test map obtained from Neurosynth (https://neurosynth.org/) using the keywords: “V1”, “MT”, “pSTS”, “A1”, “FFA”, and “place” (“PPA” did not render any results), respectively. Anatomical masks for each ROI were generated as binary coded maps (1 at the ROI, 0 everywhere else) from the Harvard-Oxford atlas downloaded from https://www.templateflow.org/. Each functional ROI was then combined by the anatomical maps to generate a combined anatomical-functional mask, from which we selected the top 200 voxels based on the z-scores within each ROI and hemisphere and thus obtained 12 ROI masks (6 ROIs*2hemispheres). To perform the ROI analyses, we averaged across the voxel-wise beta weights obtained from the whole-brain single-subject beta maps within each ROI mask for each participant to get one mean activity per hemisphere, ROI and participant. All the ROIs are shown in Supplementary Fig. S3. With the ROIs thus defined, we tested (i) whether the mean activity is significantly greater than 0 (analysis 1; one-tailed, one-sample t-test) and (ii) whether the mean activity is higher for the model fitted with socialness ratings compared to that of *chase directness* (analysis 2; one-tailed, paired t-test).

#### Optic flow

Optic flow was computed for each animation video clip using the Python package *pliers* (function *FarnebackOpticalFlowExtractor*) by computing the total amount of dense optic flow between consecutive frames (183 comparisons in total). Later, to remove an artifact (a sharp rise in optic flow in one frame every 333 ms or every 10 frames) due to an unknown reason, we identified frames whose optic flow value values that are > (median +1 SD) and replaced these timepoints with the mean values of the frames before and after it. Lastly, we normalized optic flow values across all animations (subtracted all animations’ optic flows by the minimum value and then divided each by the highest value) to the 0-1 range.

## Acknowledgements

This work was funded by R01MH129648 (E.S.F.) and Neukom CompX Faculty Grant (E.S.F.). During manuscript preparation, R.S.V. was supported by the Alexander von Humboldt Foundation. We thank Terry Sackett, Jordan Selesnick and Clara Sava-Segal for their support and guidance with setting up the fMRI data collection pipeline and the data acquisition, Ahmed Elyamani for help with developing the software, and undergraduate RAs Alison Sasaki and Ashna J. Kumar for help with quality-checking stimuli and testing the *psyanim* software.

## Declaration of generative AI and AI-assisted technologies in the manuscript preparation process

During the preparation of this work the author(s) used ChatGPT to search for research papers, identify MNI coordinates and to debug code. Each output and recommendation from ChatGPT was manually verified by the author(s). We take full responsibility for the content of the published article.

