## Supplementary material for "Subjective, not objective, socialness drives activity in the brain’s third visual pathway"

### Supplementary figures

#### Figure S1

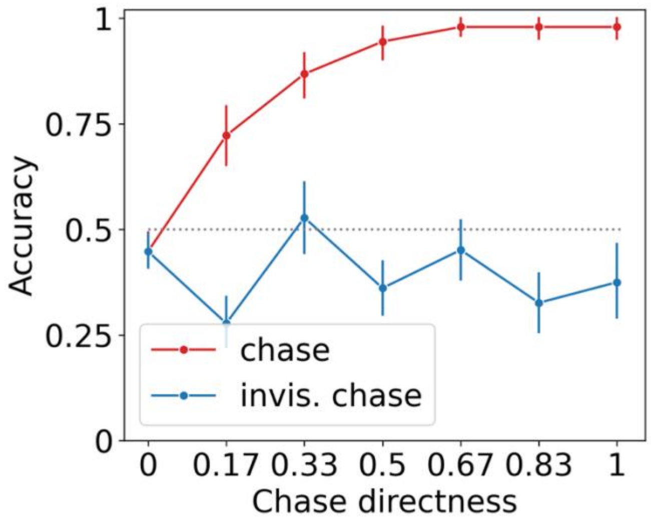


Fig S1: Group-level predator detection behavioral results from n = 24. As chase directness increases, predator detection accuracy in the true chase animations (red), but not the control animations (blue). Error bars show 95% confidence intervals across participants

### Figure S2

(a) Subjective ratings participant-level plots

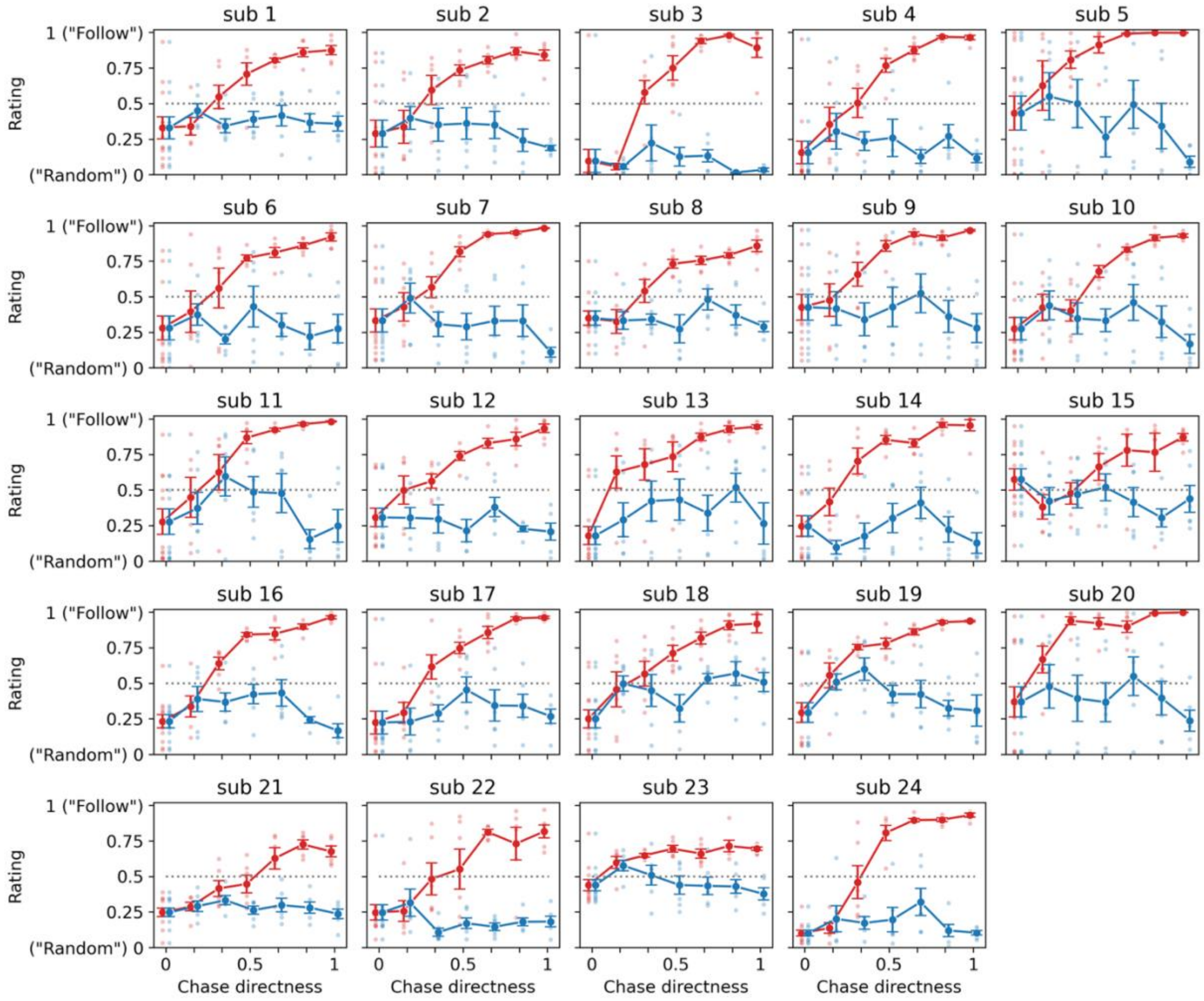

(b) Accuracy, participant-level plots

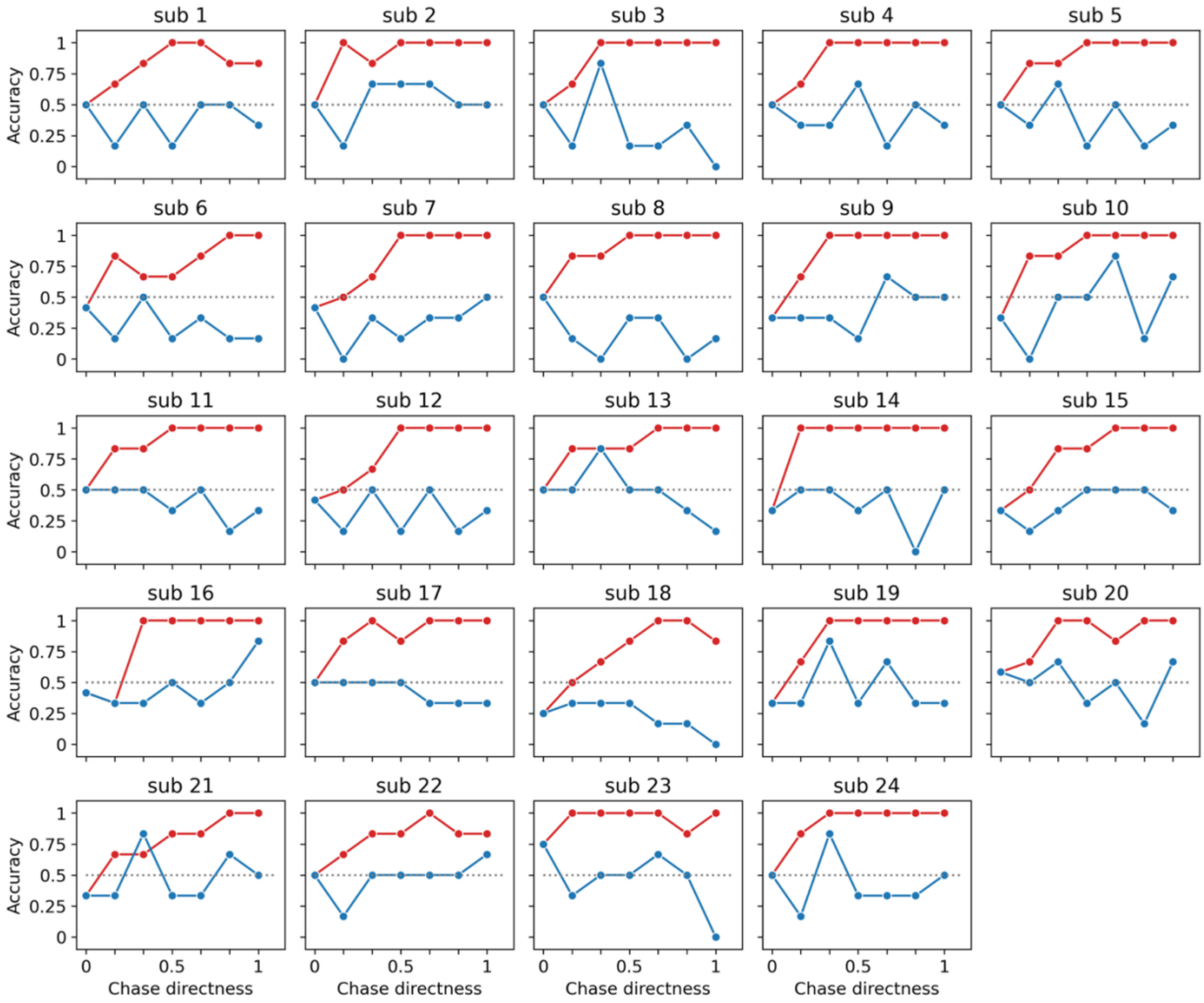


Fig S2: Participant-level behavioral data from n = 24. As chase directness increases, (a) socialness ratings and (b) predator detection accuracy increases in the true chase animations (red), but not the control animations (blue) in most participants.

Figure S3

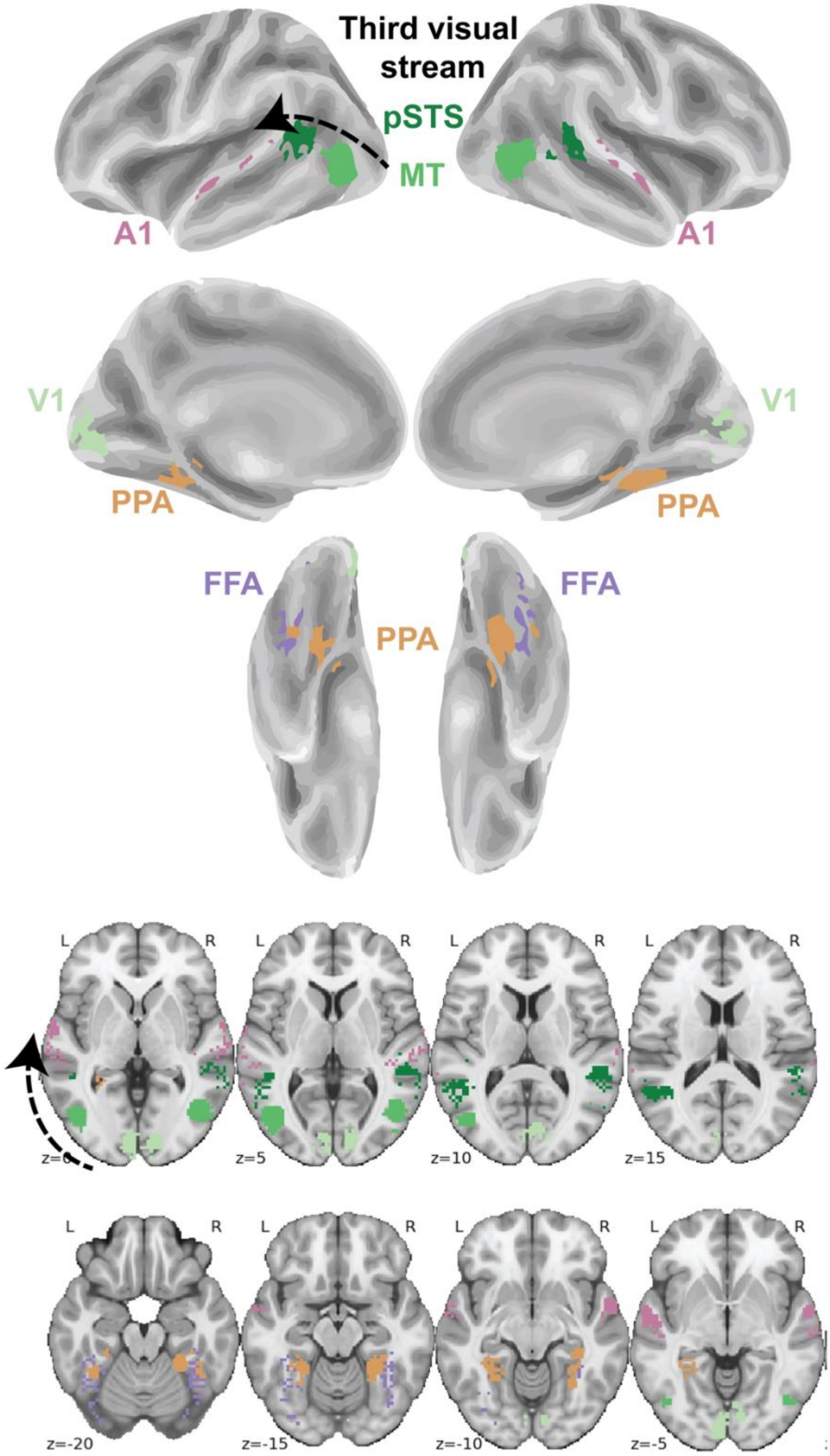
Third visual stream
pSTS
MT
A1
A1
V1
V1
PPA
PPA
FFA
FFA
PPA
L
R
z=0
z=5
z=10
z=15
z=-20
z=-15
z=-10
z=-5

Fig S3: Top 200 non-overlapping voxels within each hemisphere of each region of interest, defined by the intersection of anatomical voxels (from the Harvard-Oxford atlas) and functional voxels (from www.neurosynth.org).

### Figure S4

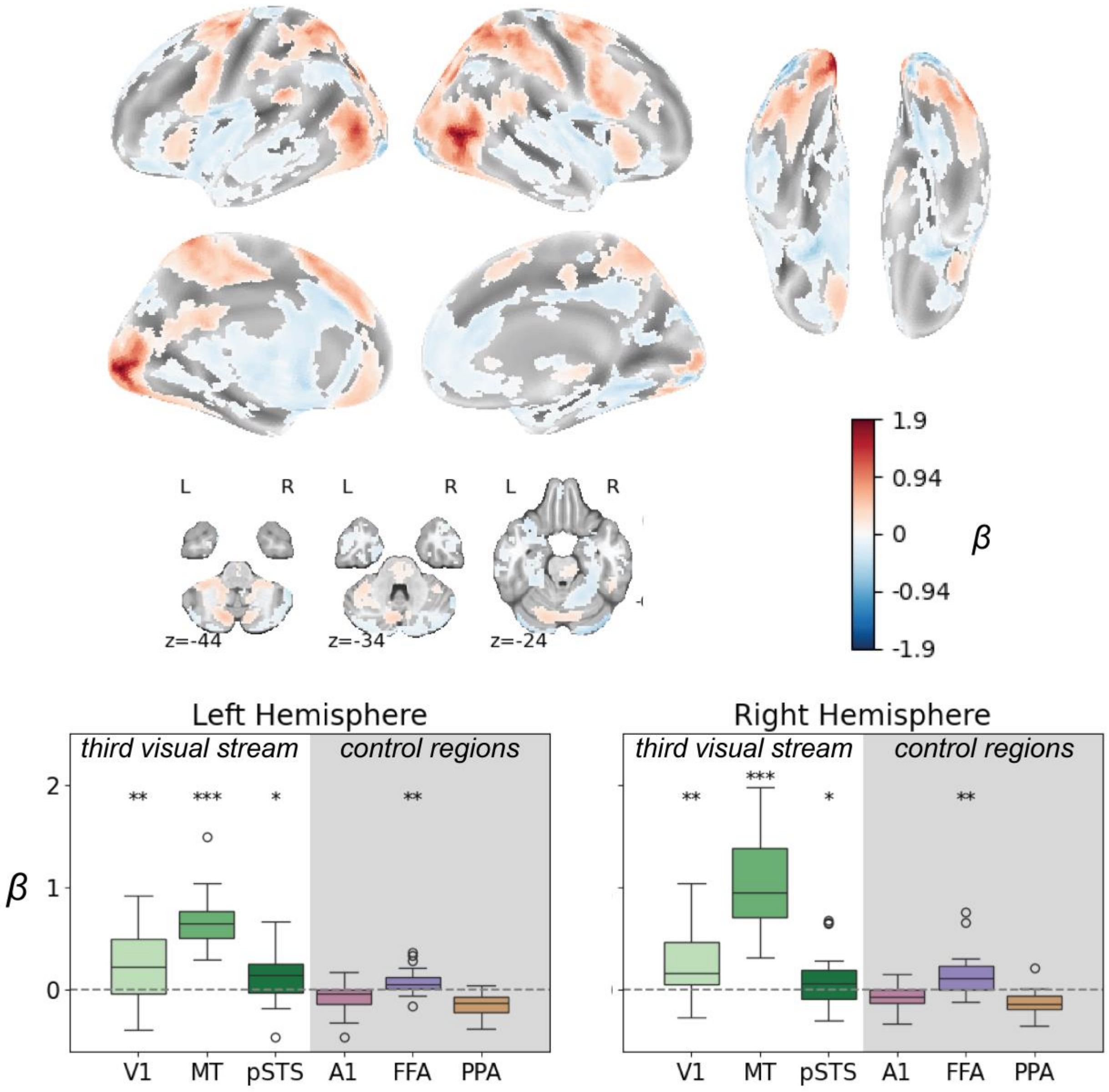


Fig S4: Neural response to optic flow in the GLM model that includes all trials and where we study the neural responsiveness to socialness (map thresholded at q < .05, corrected for multiple comparisons using the false discovery rate, plus a nominal cluster size threshold ≥ 30 voxels). The lower row shows the mean beta values across voxels within each ROI in each hemisphere. ROI definitions are shown in Supplementary Fig. S3. *** = $p < .001$, ** = $p < .01$, * = $p < .05$.

### Figure S5

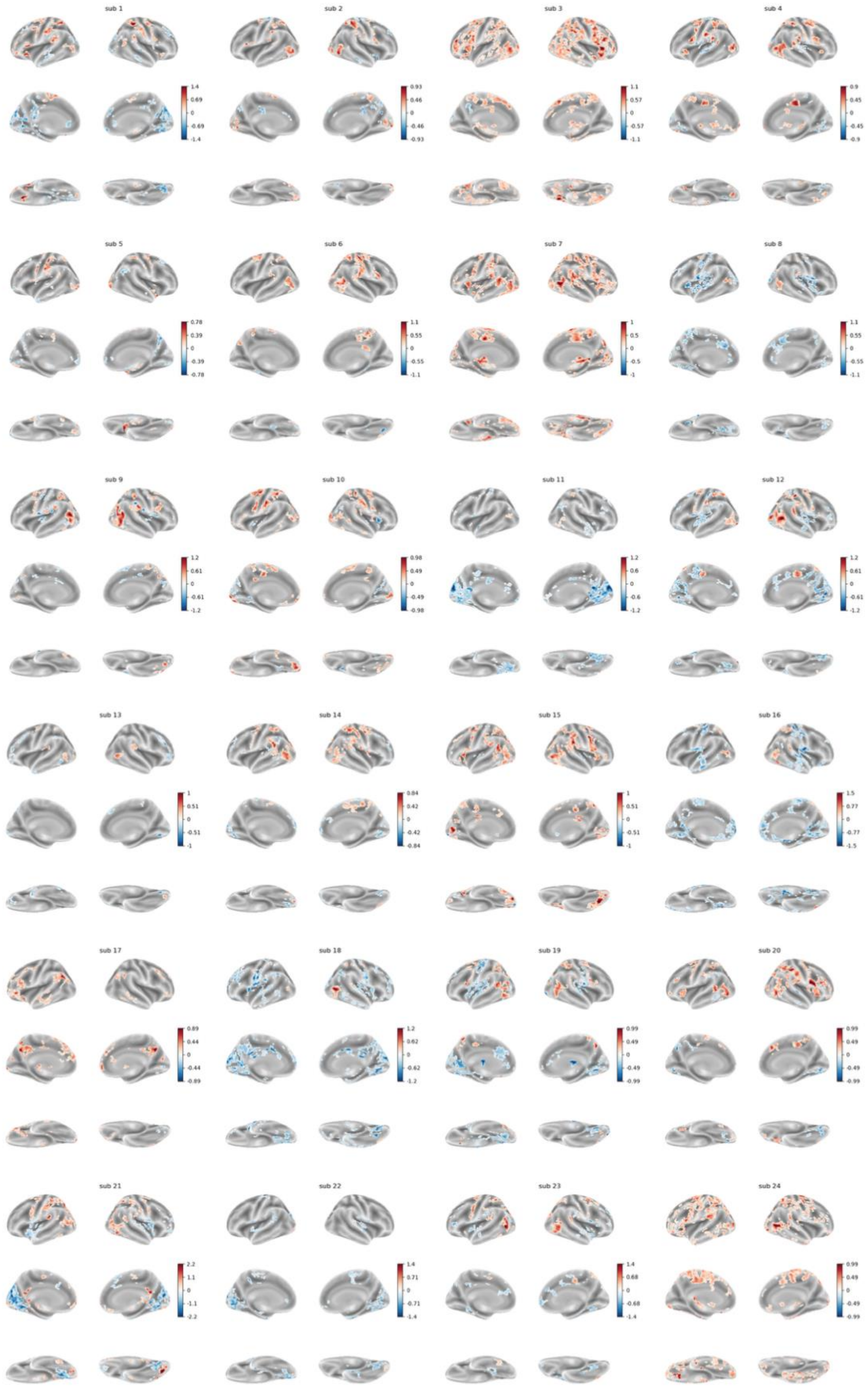
sub 1
sub 2
sub 3
sub 4
sub 5
sub 6
sub 7
sub 8
sub 9
sub 10
sub 11
sub 12
sub 13
sub 14
sub 15
sub 16
sub 17
sub 18
sub 19
sub 20
sub 21
sub 22
sub 23
sub 24

Fig S5: Participant-level brain maps of all 24 participants' BOLD responses to socialness rating (thresholded at $p < 0.05$ uncorrected for multiple comparisons, nominal cluster size >=30 voxels). These single-subject maps correspond to the group-level plot in Fig. 2.

### Figure S6

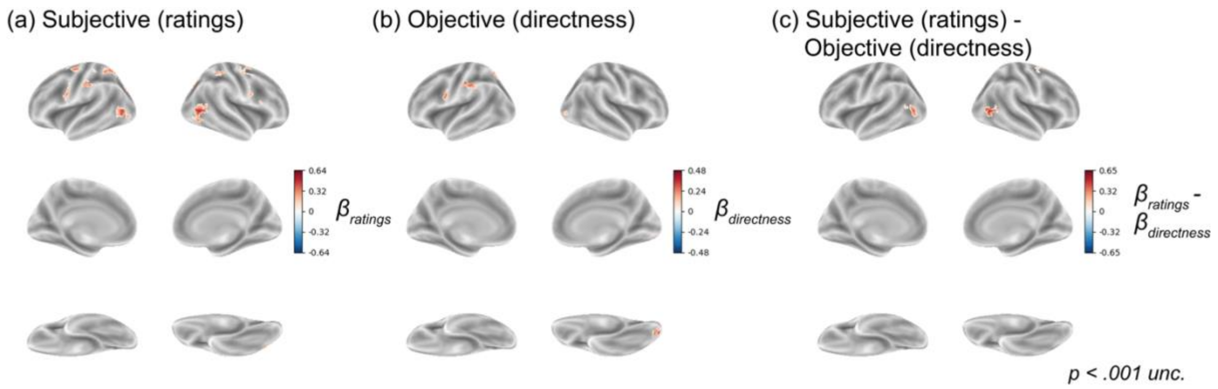


Fig S6. Complete version of main Fig. 3, including medial and ventral views. Across the brain, we only see a difference between responses to subjective ratings and objective directness in voxels around area MT in the uppermost row of all panels.

### Figure S7

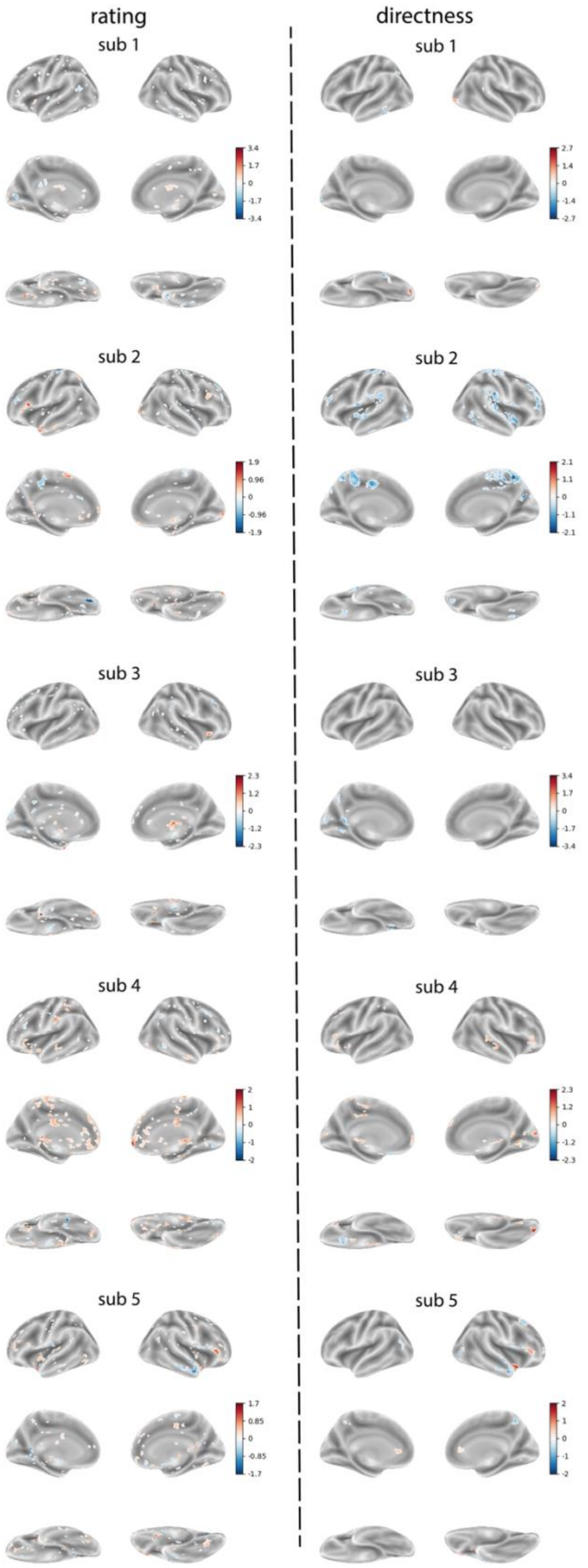
rating
directness
sub 1
sub 1
3.4
1.7
0
-1.7
-3.4
2.7
1.4
0
-1.4
-2.7
sub 2
sub 2
1.9
0.96
0
-0.96
-1.9
2.1
1.1
0
-1.1
-2.1
sub 3
sub 3
2.3
1.2
0
-1.2
-2.3
3.4
1.7
0
-1.7
-3.4
sub 4
sub 4
2
1
0
-1
-2
2.3
1.2
0
-1.2
-2.3
sub 5
sub 5
1.7
0.85
0
-0.85
-1.7
2
1
0
-1
-2

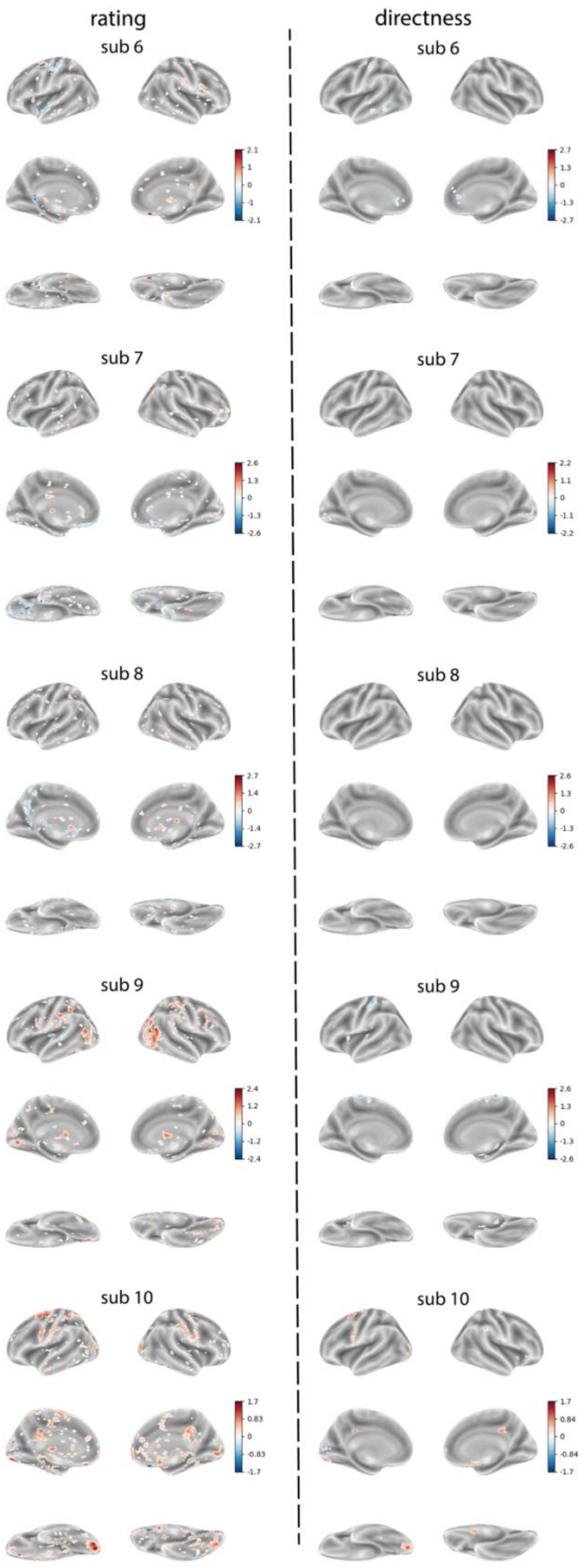
rating
directness
sub 6
sub 6
2.1
1
0
-1
-2.1
2.7
1.3
0
-1.3
-2.7
sub 7
sub 7
2.6
1.3
0
-1.3
-2.6
2.2
1.1
0
-1.1
-2.2
sub 8
sub 8
2.7
1.4
0
-1.4
-2.7
2.6
1.3
0
-1.3
-2.6
sub 9
sub 9
2.4
1.2
0
-1.2
-2.4
2.6
1.3
0
-1.3
-2.6
sub 10
sub 10
1.7
0.83
0
-0.83
-1.7
1.7
0.84
0
-0.84
-1.7

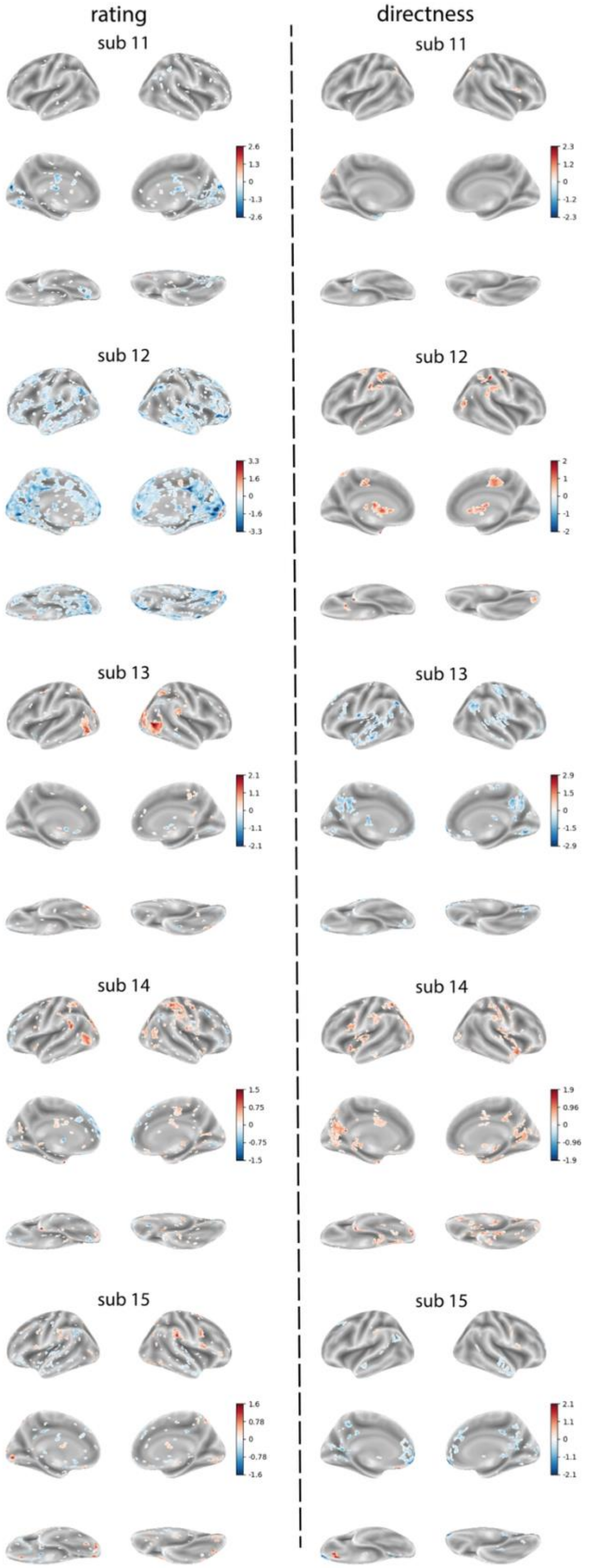
rating
directness
sub 11
sub 11
2.6
1.3
0
-1.3
-2.6
2.3
1.2
0
-1.2
-2.3
sub 12
sub 12
3.3
1.6
0
-1.6
-3.3
2
1
0
-1
-2
sub 13
sub 13
2.1
1.1
0
-1.1
-2.1
2.9
1.5
0
-1.5
-2.9
sub 14
sub 14
1.5
0.75
0
-0.75
-1.5
1.9
0.96
0
-0.96
-1.9
sub 15
sub 15
1.6
0.78
0
-0.78
-1.6
2.1
1.1
0
-1.1
-2.1

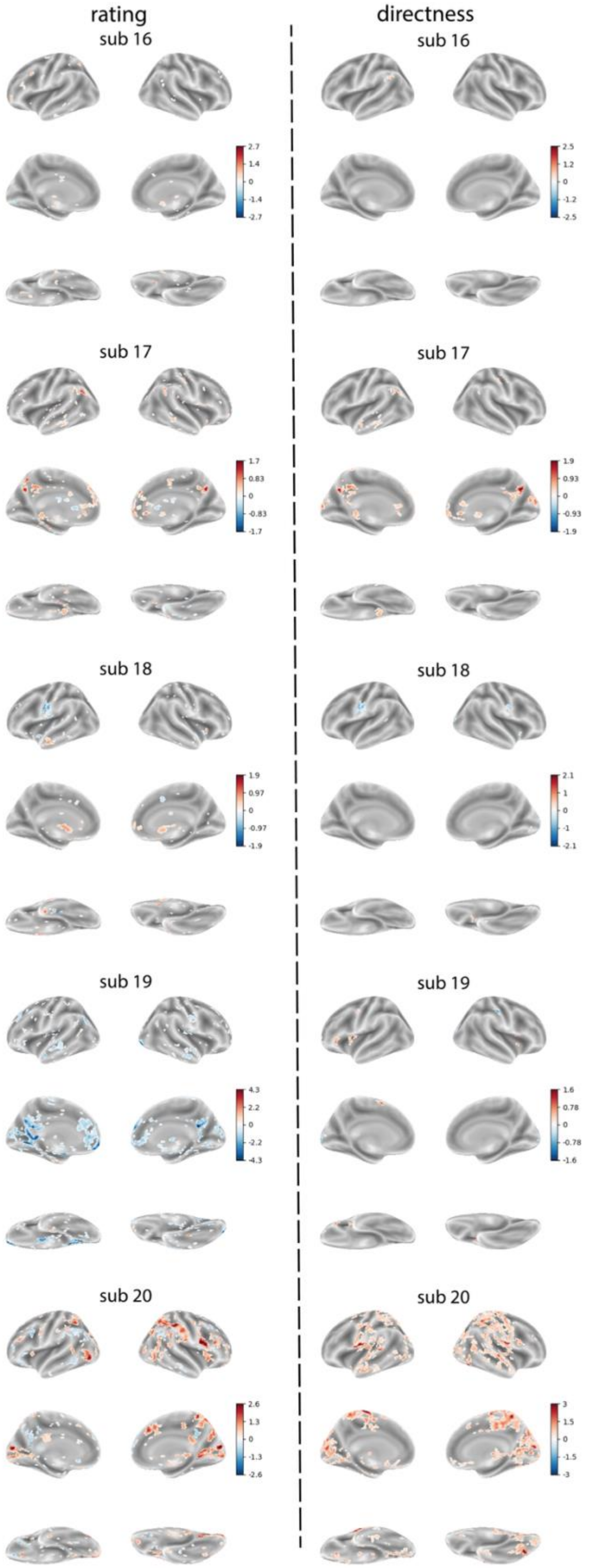
rating
directness
sub 16
2.7
1.4
0
-1.4
-2.7
sub 16
2.5
1.2
0
-1.2
-2.5
sub 17
1.7
0.83
0
-0.83
-1.7
sub 17
1.9
0.93
0
-0.93
-1.9
sub 18
1.9
0.97
0
-0.97
-1.9
sub 18
2.1
1
0
-1
-2.1
sub 19
4.3
2.2
0
-2.2
-4.3
sub 19
1.6
0.78
0
-0.78
-1.6
sub 20
2.6
1.3
0
-1.3
-2.6
sub 20
3
1.5
0
-1.5
-3

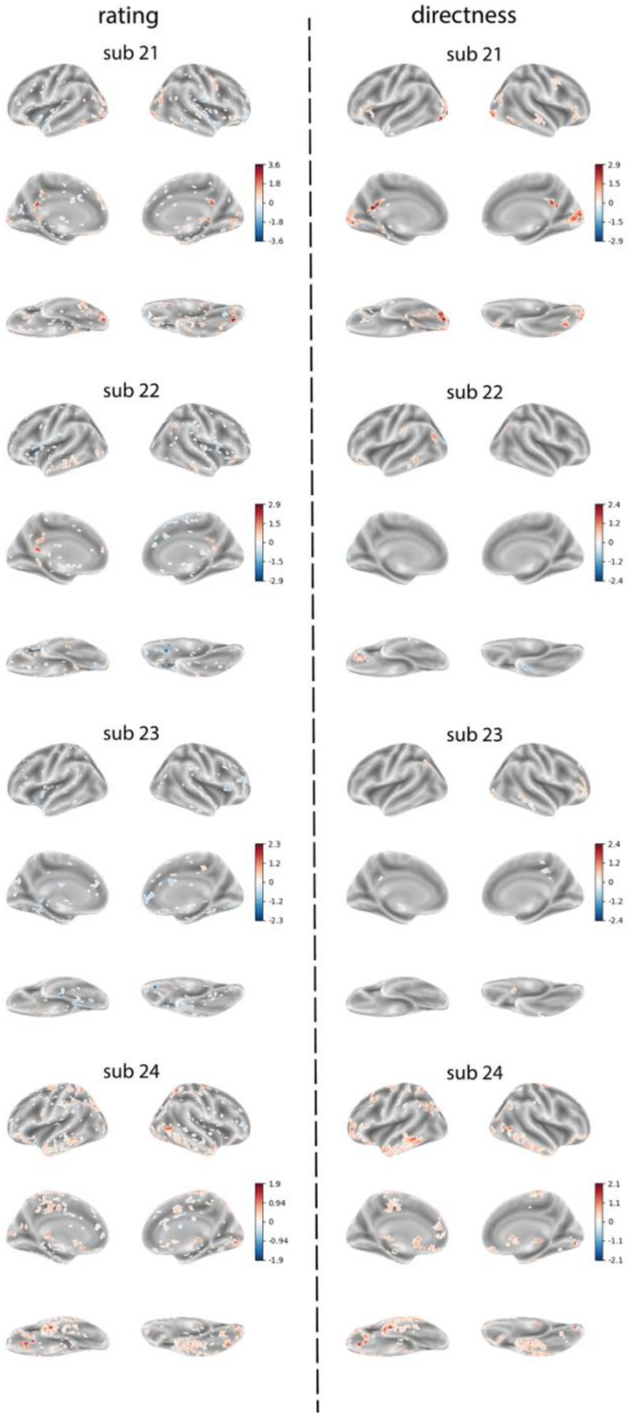


Fig S7: Participant-level brain maps from all 24 participants illustrating differences in responses to subjective ratings and objective chase directness. Each map is thresholded at $p < 0.05$ uncorrected for multiple comparisons (nominal cluster size >= 30).
